# Increased multiplexity in optical tissue clearing-based 3D immunofluorescence microscopy of the tumor microenvironment by LED photobleaching

**DOI:** 10.1101/2023.11.29.569277

**Authors:** Jingtian Zheng, Yi-Chien Wu, Evan H. Phillips, Xu Wang, Steve Seung-Young Lee

## Abstract

Optical tissue clearing and three-dimensional (3D) immunofluorescence (IF) microscopy have been transforming imaging of the complex tumor microenvironment (TME). However, current 3D IF microscopy has restricted multiplexity; only three or four cellular and non-cellular TME components can be localized in a cleared tumor tissue. Here we report a LED photobleaching method and its application for 3D multiplexed optical mapping of the TME. We built a high-power LED light irradiation device and temperature-controlled chamber for completely bleaching fluorescent signals throughout optically cleared tumor tissues without compromise of tissue and protein antigen integrity. With newly developed tissue mounting and selected region-tracking methods, we established a cyclic workflow involving IF staining, tissue clearing, 3D confocal microscopy, and LED photobleaching. By registering microscope channel images generated through three work cycles, we produced 8-plex image data from individual 400 μm-thick tumor macrosections that visualize various vascular, immune, and cancer cells in the same TME at tissue-wide and cellular levels in 3D. Our method was also validated for quantitative 3D spatial analysis of cellular remodeling in the TME after immunotherapy. These results demonstrate that our LED photobleaching system and its workflow offer a novel approach to increase the multiplexing power of 3D IF microscopy for studying tumor heterogeneity and response to therapy.

## Introduction

A solid tumor is a complex, three-dimensional (3D) ecosystem composed of malignant cancer cells and various non-malignant cells such as vascular cells, immune cells, and stromal cells^1, 2^. The multifaceted interactions of cancer cells with other normal cells lead to the development of a complicated tumor microenvironment (TME)^3^, generally presenting as highly tortuous and unevenly shaped blood vessels, dense extracellular matrix (ECM) deposition, randomly distributed immune cells, and multiple regions of hypoxia^4, 5, 6, 7^. An individual tumor has a unique TME according to the cancer type and its location in the body^8^. The diversity and spatial composition of TME components are significant determinants of cancer progression and response to therapy^9, 10^. As such, there is a considerable effort to deconvolute the TME by advancing tumor analysis techniques, which ultimately contributes to the identification of reliable TME-relevant prognostic and diagnostic biomarkers as well as therapeutic targets^11, 12^.

Recent progress in multiplex immunofluorescence (IF) has permitted the localization of multiple cellular and non-cellular TME components in a single tumor section^13, 14, 15^. Moreover, multiplex IF microscopy integrated with a fluorescence or antibody stripping method has expanded the number of TME markers visualized in a two-dimensional (2D) tumor section^16, 17^. Established multiplex imaging methods, such as t-CyCIF and Opal, have shown their successful applications in the spatial analysis of distribution and interaction of different cell types in a formalin-fixed paraffin-embedded (FFPE) or frozen tumor section^18, 19^. Nonetheless, even if fully characterized, individual tissue sections cannot adequately represent the complex 3D architecture of the TME. While 3D multiplex IF is feasible by serial cross-sectioning and 3D reconstruction of 2D tumor images^20, 21^, these tomography methods are slow and impractical for multiple tumor samples.

Optical tissue clearing and 3D optical microscopy have recently been combined as a powerful tool for high-dimensional spatial assays of various complex tissue samples^22, 23, 24, 25^. A large volumetric tumor tissue becomes optically transparent using a chemical immersion or lipid-removal method that enables matching the refractive indexes (RIs) of tumor components to either oil or water phase^26, 27^. Such optically cleared tumor tissue can be practically used for visualizing detailed 3D architecture of the TME at cellular resolution by applying various fluorescence microscopy methods, including confocal and light-sheet microscopy^28, 29, 30^. However, despite providing high dimensional image data, multiplexing capability of current 3D tissue microscopy is restricted to the number of imaging channels in a fluorescence microscope as well as the available fluorophores having different excitation and emission wavelengths, which normally allows for imaging only 3 or 4 cell markers in a 3D tumor tissue. Thus, there remains a pressing need for developing a robust approach to increase the multiplexing power of 3D tumor microscopy, enabling comprehensive spatial mapping and analysis of the TME^31^.

To address this challenge, we developed a high-power light-emitting diode (LED)-mediated photobleaching method to visualize an expanded number of TME components in 3D IF microscopy of tumor tissues. High-energetic photons emitted from a warm white LED chip at a broad range of emission spectrum (420-460 nm and 490-700 nm) can effectively and completely bleach fluorescence signals throughout an immunostained, optically transparent tumor tissue. We built a LED light-irradiating system and developed a new cyclic workflow by incorporating an LED-mediated fluorescence stripping process in the transparent tissue tomography (T3) protocol that we previously reported for 3D IF microscopy of tumor macrosections using D-fructose solution-based tissue clearing^32, 33, 34^. The cyclic workflow involves tissue processing, IF staining, optical tissue clearing, 3D fluorescence microscopy, LED photobleaching, and image data processing. A single work cycle allows us to image a maximum of four different cell markers in a whole 400 μm-thick mouse tumor macrosection. After several work cycles with LED photobleaching, we localized eight different cell markers, including cell nuclei (DAPI), vascular cells (CD31, αSMA, ER-TR7), immune cells (CD3, CD8, CD45), and cancer cells (CK8), in the same tumor macrosection using conventional 405/488/561/640 nm lasers and related emission filters in a confocal fluorescence microscope. Computational 3D image processing and analysis permitted generation of 3D multiresolution, multiplex tumor images and quantitative characterization of the 3D TME architecture. Furthermore, to test the capability of our method for spatial assessment of immunotherapy effects on the TME, we applied it to a mouse mammary tumor treated with a stimulator of interferon genes (STING) agonist and performed quantitative 3D spatial analyses of TME remodeling, immune cell infiltration, and cancer eradication at both tissue-wide and cellular resolution. Our work demonstrates that a cyclic workflow involving LED photobleaching offers a robust approach to enhance multiplexing power in 3D IF microscopy of optically cleared tumor tissues for comprehensive spatial mapping of the TME.

## Results

### LED fluorescence bleaching system for optically cleared tumor tissues

We developed a cyclic workflow for 3D multiplex IF microscopy of tumor tissues (**Figure 1A**). In the workflow, a tumor tissue is embedded in 2% agarose gel, sectioned at 400 μm thickness using a vibratome, and fixed in 2% paraformaldehyde (PFA) solution for 15 mins. Then, a tumor macrosection is subjected to repeated processes, including IF staining, optical clearing, 3D imaging, and LED-mediated fluorescence bleaching. Each work cycle produces 3 to 4 channel 3D fluorescence images using a confocal microscope. The image data are transported and saved in a data storage unit and, after completing the work cycles, collected image data are registered into a single multiplex 3D image digital file for quantitative mapping of various TME components. To effectively strip fluorescence signals in tumor macrosections, we built a photobleaching system using a high-power LED chip (warm white, 100W) and cooling units (**Figure 1B**). A tumor macrosection mounted on a 1 mm-thick glass slide was placed on the sample stage at a 2 cm distance from the LED chip surface. As the high-power LED chip generates heat when its light is on, a heatsink and fan were assembled to the bottom of the LED chip for cooling the photobleaching device. Also, to prevent heat-induced damage of tissue samples during light irradiation, we placed the LED device in a refrigerator containing dry ice, which maintained the temperature on the top of LED chip between 1 and 5°C over 8 hrs of light irradiation (**Figure 1C**). We also confirmed the 100W warm white LED chip produces bright light (8 - 9K Luminous Flux) with broad emission bands (420-460 nm and 490-700 nm), which covers the 488, 561, and 633 nm excitation wavelengths of widely used fluorophores, such as FITC, Cy3, Cy5 and DyLight (DL) 488, 550, 633 dyes (**Figure 1D**).

**Figure 1.**
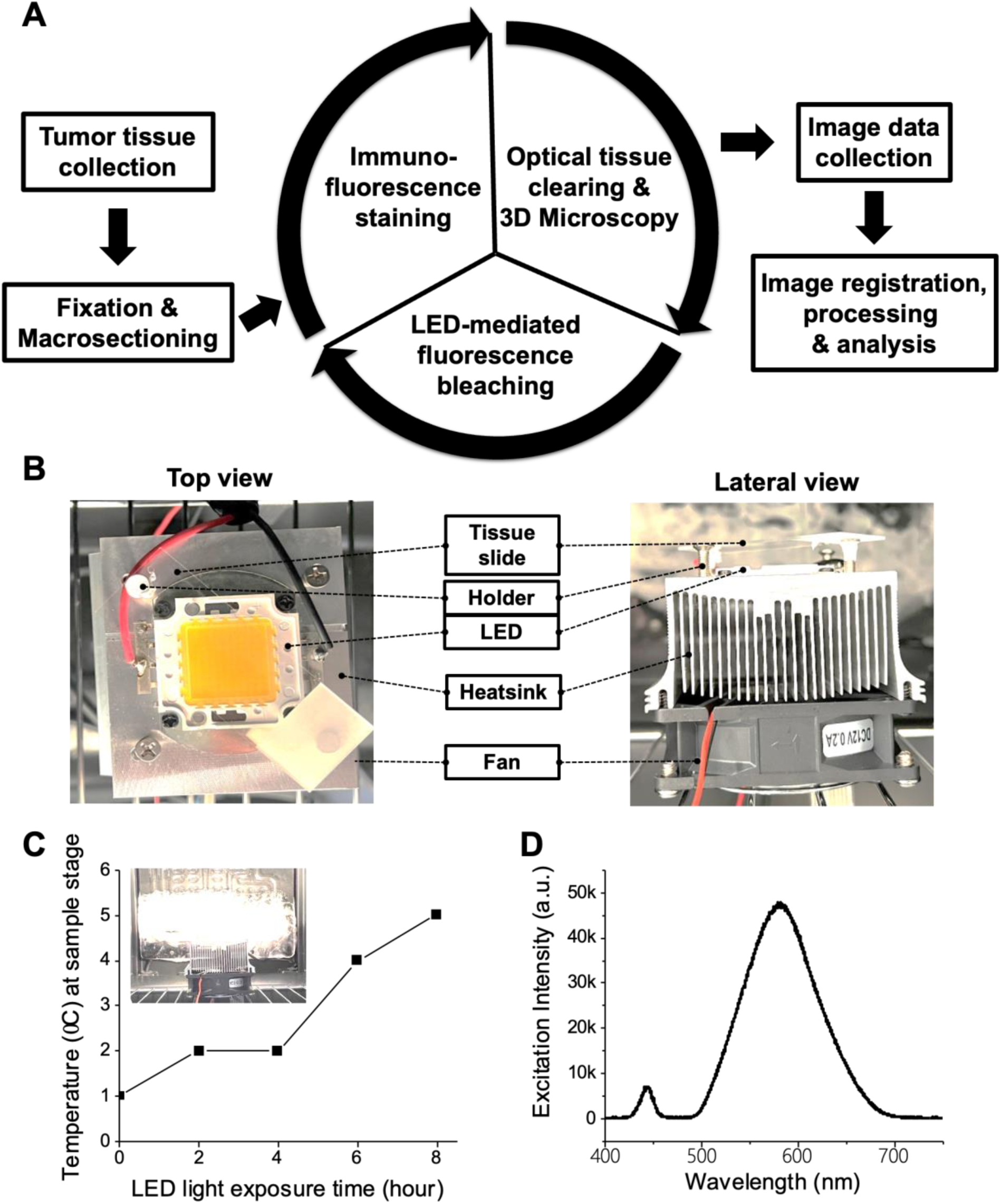
High-power LED irradiation system for 3D multiplex immunofluorescence microscopy of optically cleared tissues. **(A)** Cyclic workflow involving tissue processing, immunofluorescence staining, optical tissue clearing, 3D optical microscopy, LED light irradiation, and computational image processing. **(B)** Photographs of high-power LED light irradiation system assembled with cooling units (**left**: Top view, **right**: Lateral view). **(C)** Temperature change at the sample stage of the LED system during light irradiation (0 - 8 hours). The inset photograph shows the ‘light-on’ LED system in a refrigerator. (**D**) Emission spectrum of 100W Warm White LED chip in the system.

To test 3D fluorescence bleaching using the LED system, we prepared a 400 μm-thick macrosection from a mouse mammary TUBO tumor using a vibratome and stained with a cocktail of fluorescent primary antibodies such as anti-ER-TR7-FITC, αSMA-Cy3, and CD31-Cy5. The tumor macrosection was then incubated in a series of D-fructose solutions for optical tissue clearing. Our previous reports showed that simple immersion of tumor macrosections in high concentration D-fructose solutions enabled the achievement of adequate tissue transparency for 3D confocal fluorescence microscopy^32, 33^. After tissue clearing, we loaded the macrosection on the LED system and exposed it to warm white light for 8 hrs. Using a confocal microscope, we monitored changes of the fluorescence intensities of FITC, Cy3, and Cy5 dyes in the 488, 561, and 633 nm channels, respectively (**Figure 2A, B**). The tissue-wide 2D images were obtained from the bottom (LED-close side) and top (LED-far side) surfaces of the tumor macrosection at 0, 2, 4, and 8 hrs post LED light exposure. All fluorescence signals on the both sides of the macrosection were dramatically decreased to around 14-17% of the initial intensities after the first 2 hrs of exposure to LED light, and further declined and completely bleached after an additional 6 hrs of irradiation. This result shows that a high-power warm white LED system effectively removes fluorescence signals over a broad wavelength range and throughout a transparent tumor macrosection.

**Figure 2.**
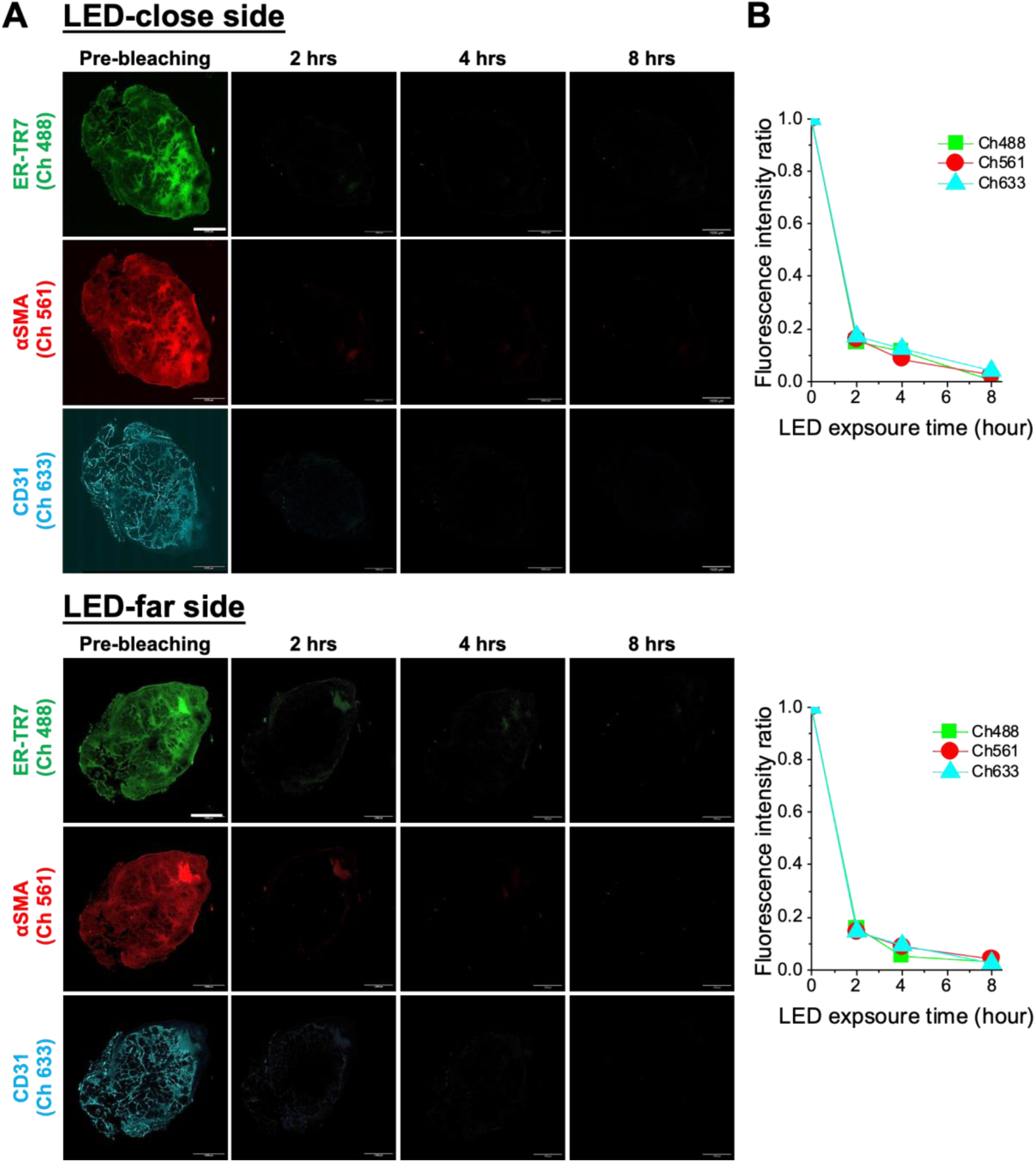
LED-mediated fluorescence bleaching in a tumor macrosection. **(A)** Mosaic fluorescence channel images of ER-TR7 (green, excitation at 488 nm), αSMA (red, excitation at 561 nm), CD31 (cyan, excitation at 633 nm) on the LED-close side (bottom surface) and -far side (top surface) of a 400 μm-thick tumor macrosection before and after exposure to warm white LED light for 2, 4, and 8 hours. Scale bar: 1 mm. **(B)** Quantification of fluorescence intensity changes in 488, 561, and 633 channel images from the LED-close side and -far side of the tumor macrosection during LED-mediated photobleaching. The intensity ratio was plotted based on the pre-IF staining ‘0’ and pre-bleaching ‘1’ levels.

We additionally investigated the feasibility of color light LED, which has a narrow emission band, for selective photobleaching of a target fluorescence in an IF-stained 3D tumor tissue. We measured the spectrum of blue, cyan, green, yellow, and red light from different 100 W LED chips (**Supplementary Figure 1A**), and prepared green and red light LED photobleaching systems in the same way that the warm white LED system was built (**Supplementary Figure 1B**). Green and red color light from the LED systems showed narrow emission spectra of 484-587 nm (peak: 526 nm) and 586-678 nm (peak: 642 nm), respectively. After immunostaining with anti-αSMA-DyLight (DL) 488, ER-TR7-Dylight (DL) 550, and CD31-Cy5 antibodies and optically clearing in D-fructose solutions, 400 μm-thick TUBO tumor macrosections were exposed to green and red color light for 12 hrs. We measured the fluorescence intensities on the top surfaces (LED-far side) of the macrosections at 0, 2, 4, 8, and 12 hrs post-light irradiation (**Supplementary Figure 2A, B).** Exposing to green light significantly reduced the signals of both DL488 and DL550 by up to 89% after the first 2 hrs and completely bleached at 12 hrs, while the signal of Cy5 slowly faded during exposure to green light and remained more than 62% of the initial intensity at 12 hrs post-irradiation. For the macrosection exposed to red LED light, a fast and large reduction (up to 83%) of the intensity of Cy5 was observed after the first 2 hrs of photobleaching, but the fluorescence signals of both DL488 and DL550 gradually declined over 12 hrs exposure. Although color light from the high-power LED chip has some bleaching effect to all the fluorophores having different excitation wavelengths, the results demonstrate the potential of color light LED systems as a tool for selectively bleaching a target fluorescence in a 3D tumor tissue. Despite the need for further optimization in selection of photo-stable and -unstable fluorophores and adaptation of optical narrow bandpass filters, color light LED systems might be useful for direct *in situ* analysis of the distribution and interactions of multiple cellular components along with a spatial landmark in the TME. For example, one could selectively bleach and switch fluorescent antibodies for visualizing various cellular makers in hypoxic areas while maintaining the fluorescence signal of the reference landmark.

### LED photobleaching-based 3D multiplex immunofluorescence microscopy

We implemented the cyclic workflow with fluorescent antibodies to image panels of vascular cells, immune cells, and cancer cells in a tumor macrosection (**Table 1** **and** **Figure 3A**). A TUBO tumor was cast in 2% agarose gel and a 400 μm-thick macrosection was cut using a vibratome. The gel surrounding the macrosection was marked for orientation on microscopy to facilitate X-Y alignment in the registration of image data (**Supplementary Figure 3A**). The tumor macrosection was stained with anti-ER-TR7-FITC, αSMA-Cy3, and CD31-Cy5 antibodies for visualizing vascular fibroblast, smooth muscle cells, and endothelial cells in the TME, respectively. After optically cleared in D-fructose solution, the macrosection was mounted on a glass slide with a frame and covered with a glass slip on the top. We could find and start 3D scanning of the macrosection from the same Z position in each work cycle by using the 785 nm excitation channel and open emission filter on an upright confocal fluorescence microscope (Caliber ID, RS-G4) that provided a reflection image originating from the tumor tissue (**Supplementary Figure 3B**). After 3D imaging of the vascular cells in the whole tumor macrosection on three fluorescence excitation and emission channels (488, 561, 633 nm), the macrosection was placed in the LED photobleaching system and exposed to warm white light for 8 hrs to remove all existing fluorescence signals. The macrosection was then washed in PBS, stained with anti-CD8-DL488, CD45-DL550, CD3-DL633 antibodies for immune cells, optically cleared in D-fructose solutions, and imaged in the same way as described above. After LED photobleaching, cytokeratin (CK) 8-postive cancer cells were detected after repeating the workflow. Collected fluorescence channel images were registered to reconstruct a 3D multiplex image using Fiji (ImageJ) plugins (**Figure 3B**). A 7-plex 3D image of vascular cells, immune cells, and cancer cells in the TME of a whole TUBO tumor macrosection was achieved using LED photobleaching system and cyclic workflow (**Supplementary Video 1**).

**Figure 3.**
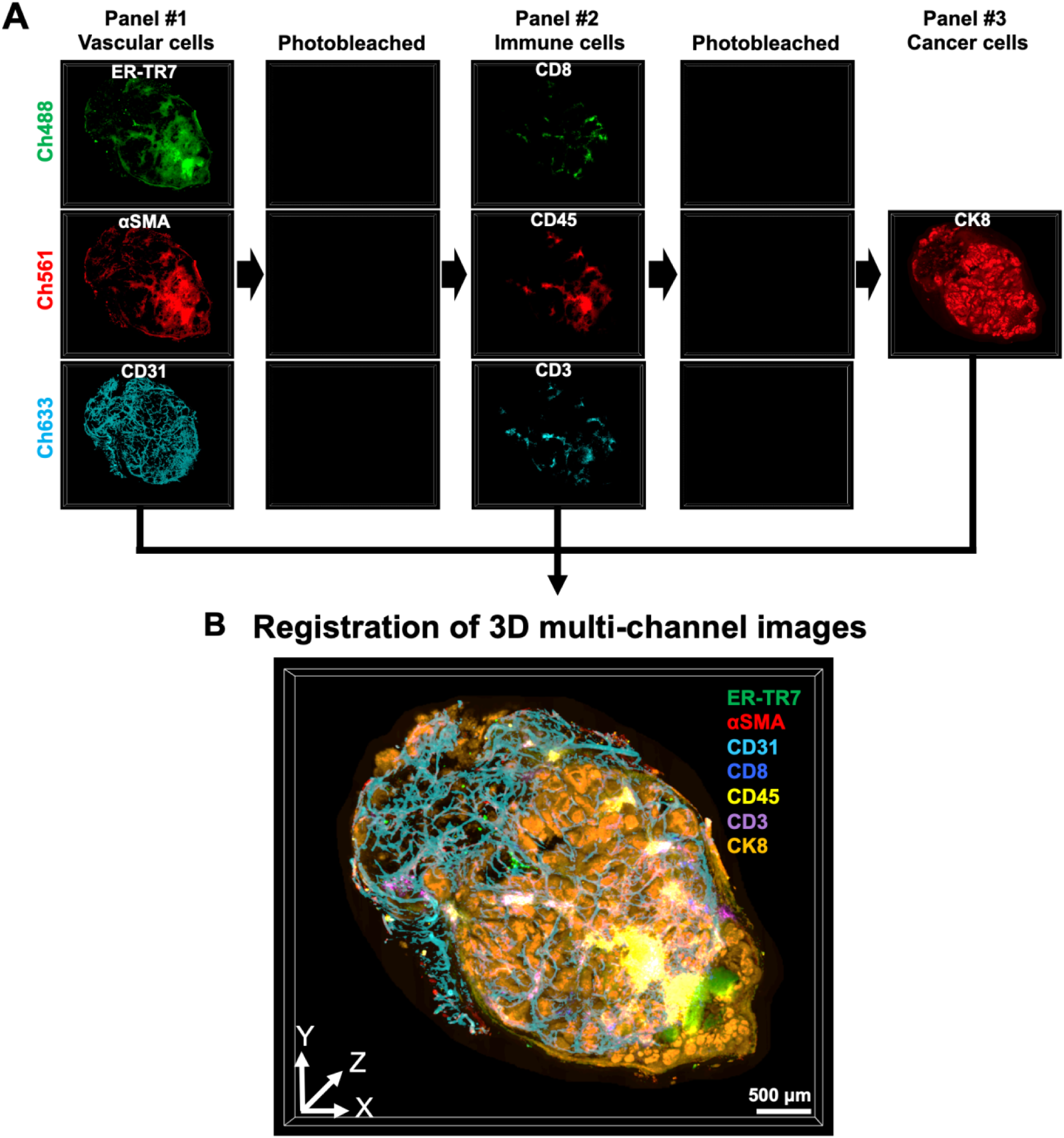
Flow of 3D multiplex immunofluorescence microscopy for visualizing multiple cell types in a whole tumor macrosection. **(A)** Repeated ‘Staining-Imaging-Photobleaching’ process using the warm white LED light irradiation system. Markers for vascular cells (ER-TR7, αSMA, CD31), immune cells (CD8, CD45, CD3), and cancer cells (CK8) were sequentially localized in the 488, 561, and 633 channel images of a 400 μm-thick tumor macrosection through three work cycles (**Table 1**). **(B)** 3D rendering of reconstructed 7-plex image of the cell markers in the tumor macrosection after computational alignment and registration of the multiple channel images. Scale bar: 500 μm.

**Table 1.** Fluorescence antibody panels for validating the cyclic workflow of 3D multiplex immunofluorescence microscopy with a tumor macrosection.

| Channel/Panel | #1- Vascular cells | #2- Immune cells | #3- Cancer cells |
| --- | --- | --- | --- |
| Ch488 | ER-TR7-FITC | CD8-DL488 |  |
| Ch561 | αSMA-Cy3 | CD45-DL550 | CK8-DL550 |
| Ch633 | CD31-CY5 | CD3-DL633 |  |

We further developed a registration method for cellular resolution 3D image data *via* DAPI-mediated cell nuclei staining (**Supplementary Figure 4**). We stained a 400 μm-thick TUBO tumor macrosection with DAPI, optically cleared in D-fructose solution, and exposed to warm white light using the LED photobleaching system. Different from other fluorophores having a long-wavelength absorption band, the intensity of DAPI (excited at shorter wavelengths between 300 nm and 418 nm) was slightly decreased to 83% after 16 hrs photobleaching (**Supplementary Figure 4A, B).** Using the signal pattern in the DAPI channel as a spatial reference, we could repeatedly identify and image the same selected region within a tumor macrosection using a 40x objective in 3D over three work cycles, which permitted the alignment and reconstruction of high-resolution 3D multiplex images (**Supplementary Figure 4C**). These results show that the cyclic workflow with LED photobleaching offers multiresolution, multiplex 3D microscopy images of tumor tissues for robust spatial mapping of the TME.

### Multiplex 3D spatial analysis of immunotherapy effects on the tumor microenvironment

To validate our method as a tool for evaluation of cancer immunotherapy, we performed multiresolution 3D spatial mapping of various cellular TME components in a TUBO tumor treated with a STING agonist, 5,6-dimethylxanthenone-4-acetic acid (DMXAA). Compared to a control untreated tumor, we quantitatively determined the structural remodeling of the TME in a TUBO tumor after local treatment with DMXAA (**Figure 4**). A single 400 μm-thick macrosection was selected from the middle of each control and treated TUBO tumor and underwent the cyclic workflow with DAPI and the listed fluorescent antibodies (**Table 2**) for 3D multiplex microscopy of diverse cell types at tissue-wide and cellular resolution. Constructed 3D multiplex images of the tumor macrosections displayed detailed architecture of the TME composed of cancer cells (CK8), vascular cells (αSMA, CD31, ER-TR7), and immune cells (CD8, CD45, CD3). DAPI signal represents the locations of all cellular components in the TME. The vascular and cancer cells that mainly structured the TME were segmented in the tumor macrosection images using Fiji (ImageJ) software (**Supplementary Figure 5**). It is clearly noticed in the whole macrosection images that CK8^+^ cancer cells broadly localized and overall occupied more than 68% of the parenchyma in the control tumor, while CK8^+^ cancer cells were unevenly distributed and occupied less than 30% of the cellular area of the treated tumor (**Figure 4A, B**). We also found the relative volumes of αSMA^+^ vascular smooth muscle cells (0.1%), CD31^+^ endothelial cells (8%), and ER-TR7^+^ fibroblasts (32%) in the treated tumor were lower than those in the control tumor (2%, 26%, and 49% of αSMA^+^, CD31^+^ and ER-TR7^+^ cells, respectively), corresponding to previous reports that showed DMXAA-mediated tumor vasculature disruption triggered by a strong immune response^35, 36^. In addition, compared to the treated tumor, larger volumes of thicker and denser segments of ER-TR7^+^ fibroblast-abundant stroma were observed in the control tumor, especially in the center area (**Supplementary Figure 6 and** **Figure 4C**). High-resolution 3D images display distinct patterns of regional ER-TR7^+^ fibrotic stroma between the control and treated tumors in detail (**Figure 4A** (bottom rows of each tumor) and **Supplementary Figures 7, 8**). We noticed in the high-resolution 3D images that large numbers of individual CD8^+^, CD45^+^, CD3^+^ immune cell types were localized in ER-TR7^+^ stromal areas of the control tumor (**Figure 4D**).

**Figure 4.**
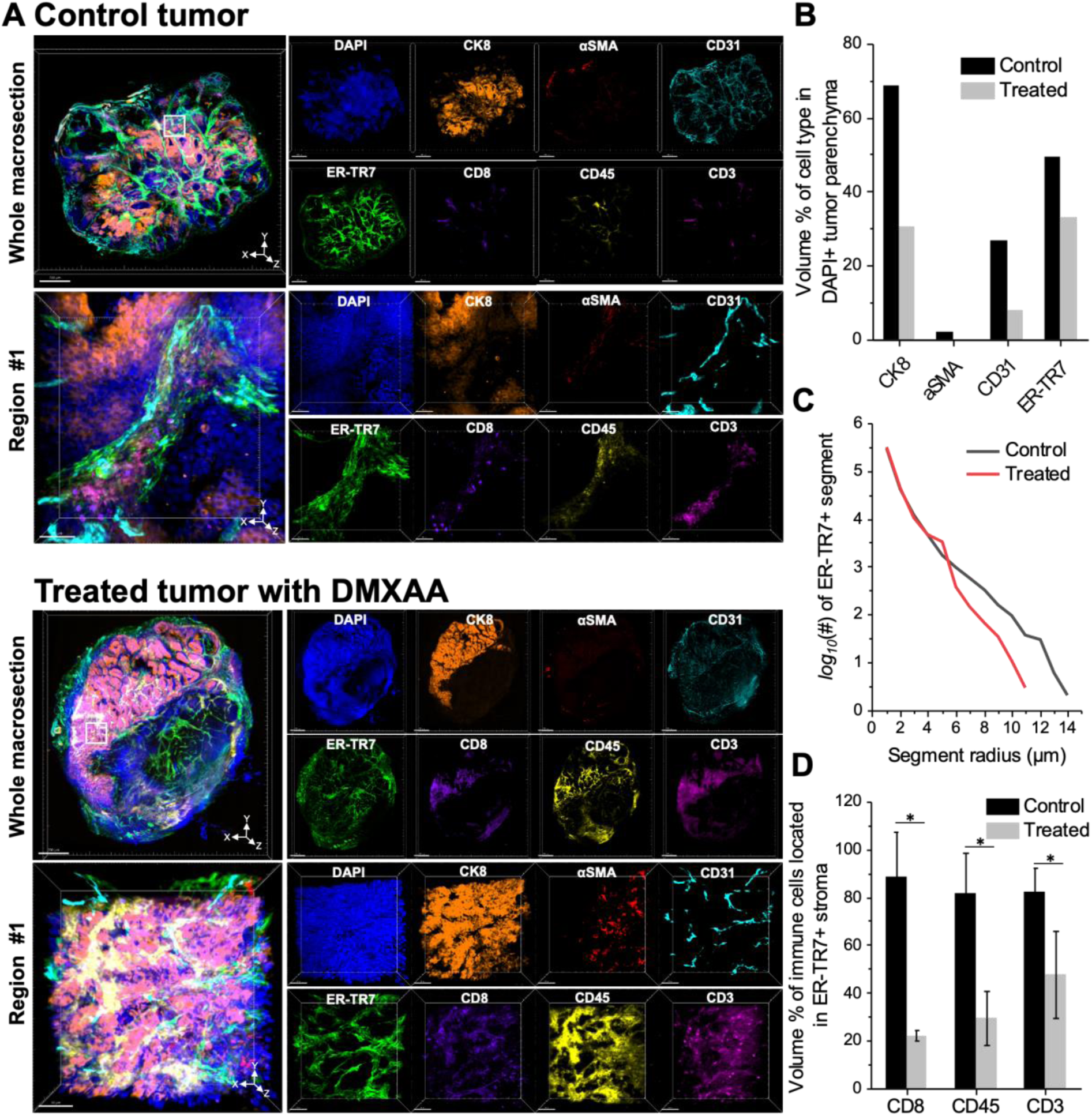
Quantitative 3D spatial analysis of immunotherapy effect on multiple cell types in the tumor microenvironment. **(A)** 3D images of vascular cells (ER-TR7, αSMA, CD31), immune cells (CD8, CD45, CD3), and cancer cells (CK8) in macrosections from a control non-treated tumor and treated tumor with DMXAA at whole-tissue (top rows) and cellular (bottom rows) levels. Scale bars: 700 μm and 50 μm. High-resolution 3D images of Region #1 were achieved from the white boxes in the control and treated tumor macrosections. **(B)** Volume percentages (%) of ER-TR7^+^, αSMA^+^, CD31^+^ and CK8^+^ segments within the DAPI^+^ parenchyma of the whole control and treated tumor macrosections. **(C)** Distribution of volume (%) of ER-TR7^+^ fibrotic stroma segments in the whole control and treated tumor macrosections according to the radius of ER-TR7^+^ segments. **(D)** Volume percentages (%) of CD8^+^, CD45^+^, and CD3^+^ immune cells located in ER-TR7^+^ fibrotic stroma segments in four different regions (additional Regions #2-4 in **Supplementary Figure 7, 8**) of the whole control and treated tumor macrosections. Values are displayed as mean ± SD, and statistics were run using multiple unpaired t-tests (*\*P* < 0.01).

**Table 2.** Fluorescence antibody panels for 3D spatial analysis of immunotherapy effects on the tumor microenvironment.

| Channel/Panel | #1- Vascular cells | #2- Immune cells | #3- Cancer cells |
| --- | --- | --- | --- |
| Ch405 | DAPI | DAPI | DAPI |
| Ch488 | αSMA-DL488 | CD8-DL488 | |
| Ch561 | ER-TR7-DL550 | CD45-DL550 | CK8-DL550 |
| Ch633 | CD31-CY5 | CD3-DL633 |  |

Next, we further investigated the immune cell subtypes and their interactions with cancer cells by quantitatively analyzing the high-resolution 3D image of a representative region, such as region #1 of the control and treated tumors where all immune cell markers and CK8^+^ cancer cells are captured (**Figure 5**, **Supplementary Figure 9, and Videos 2, 3**). We first extracted individual CD8^+^, CD45^+^, CD3^+^ immune cell types and further classified into four different CD45^+^ immune cell subtypes: CD45^+^CD8^-^CD3^-^, CD45^+^CD8^+^CD3^-^, CD45^+^CD8^-^CD3^+^, and CD45^+^CD8^+^CD3^+^ cells (**Figure 5A**). Based on the immune cell segments, we measured the volume percentages (%) of CD45^+^ immune cell subpopulations in region #1 of the control and treated tumors (**Figure 5B**). While the larger volume % of non-T lymphocyte population (CD45^+^CD8^-^CD3^-^) was detected in region #1 of the control tumor, the larger volume % of cytotoxic T lymphocytes (CD45^+^CD8^+^CD3^+^ cells, CTLs) was observed in region #1 of the treated tumor. We further interrogated the spatial distribution of CD45^+^CD8^+^CD3^+^ CTLs, the main effector cells required for effective cancer immunotherapy (**Figure 5C**)^37, 38^. More than 22% of CK8^+^ cancer cells physically contacted CTLs in region #1 of the treated tumor, while only around 2% of CK8^+^ cancer cells interacted with CTLs in region #1 of the control tumor (**Figure 5D**). The results suggest the treatment with DMAXX increased the tumor infiltration of CTLs and eradicated cancer cells by a CTL-mediated immune response.

**Figure 5.**
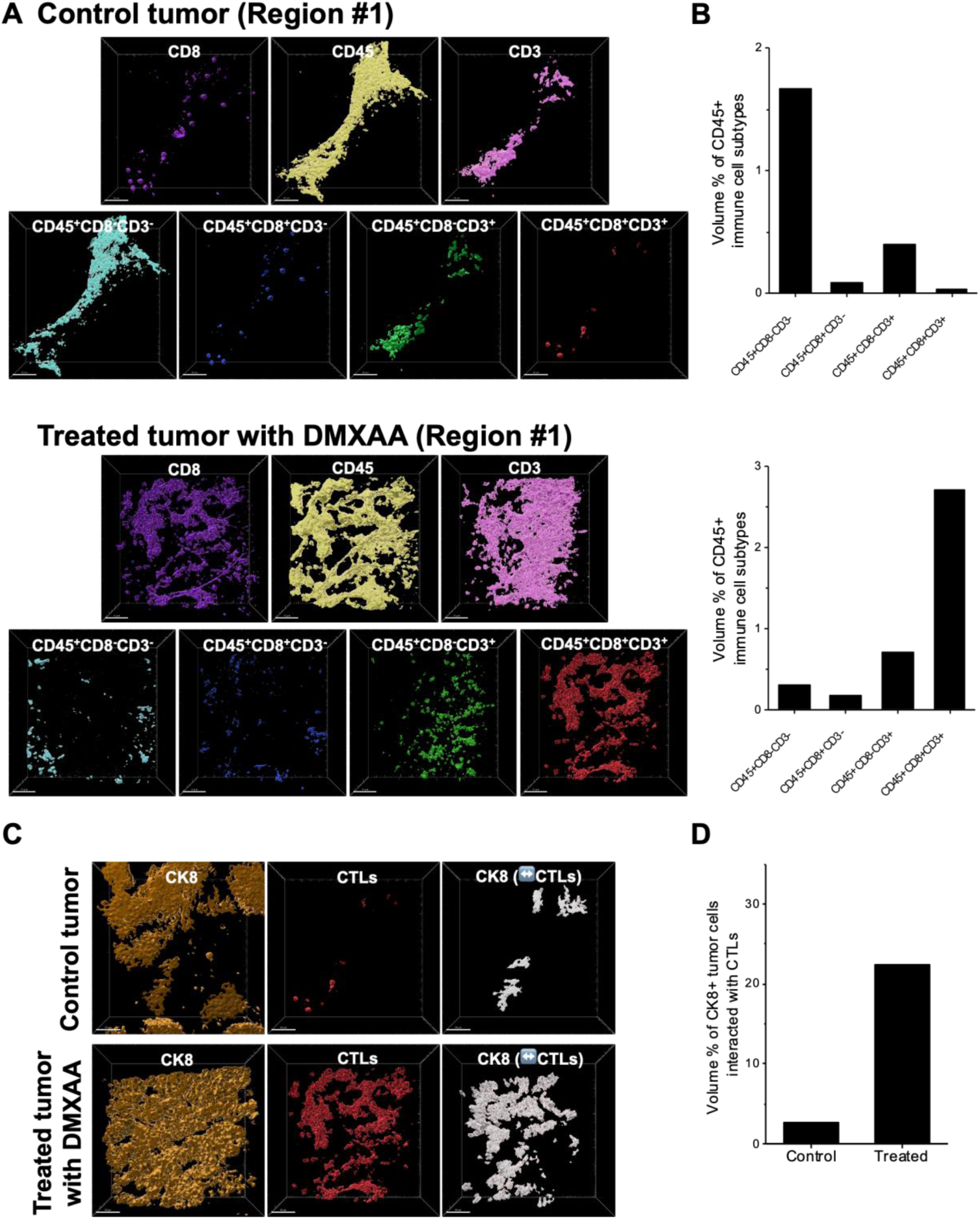
Quantitative 3D spatial analysis of tumor-infiltrating immune cell subtypes. **(A)** Segmentation of CD45^+^ immune cell subtypes in Region #1 of control and treated tumor macrosections according to CD8- and CD3-negative and -positive expression. **(B)** Volume percentages (%) of CD45^+^ cell subtypes segments in each Region #1. **(C)** Segmentation of CK8^+^ tumor cells in contact with CD45^+^CD8^+^CD3^+^ cytotoxic T lymphocytes (CTLs) in Region #1 of control and treated tumor macrosections. **(D)** Volume percentages (%) of CK8^+^ tumor cell segments interacting with CTLs over the total CK8^+^ area in each Region #1.

## Discussion

Over the past few years, optical tissue clearing technology has been significantly evolving and transformed optical microscopy of tissues in broad fields of research including histopathology, neuroscience, and development biology^39, 40, 41^. Various immersion methods using aqueous and organic solvents yield a high transparency in large volumetric tissues (e.g. whole mouse brain and human organs) with preservation of cellular and molecular integrity in the tissues^42, 43^. Together with advanced fluorescence microscopy and IF techniques, optical tissue clearing enables 3D IF microscopy of preclinical and clinical tumor tissues, offering unprecedented 3D views of the spatial architecture of the TME at cellular resolution^44^ ^45, 46^. This integration has opened a new field of ‘3D digital histopathology’. This field takes advantage of two powerful tools: 1) 3D tumor microscopy, which provides 3D landscape of the TME, and 2) artificial intelligence (AI) and machine learning-based computational image data analysis methods^47^, which assist pathologists for precise cancer diagnosis and prognosis. However, despite producing high-dimensional cell-resolution image data, current 3D IF microscopy methods can image only 3 or 4 cell markers in a tumor tissue, which limits its applications for in-depth investigation of the structural organization and spatial interactions of diverse cell types in tumor. To better understand the TME, it is necessary to address the bottleneck of multiplexing capability in 3D IF microscopy.

Fluorescence and antibody stripping methods have been widely used for multiplex two-dimensional (2D) imaging of tumor sections less than 10 μm thick^48, 49^. For tumor sections stained with fluorescent primary and secondary antibodies, treatment with hydrogen peroxide (H_2_O_2_) oxidizes fluorophores and eliminates fluorescence signals on the tissue sections^16^. It enables multiplex 2D IF imaging of tumor sections by looping three processes (IF staining, imaging, and fluorescence stripping)^18, 50^. For tumor sections stained with horseradish peroxidase (HRP)-conjugated antibodies and HRP-reactive fluorophores (e.g. tyramide signal amplification (TSA) dyes), implementing heat-induced antigen retrieval (HIAR) degrades HRP-antibodies bound to a protein antigen on a formalin-fixed, paraffin-embedded (FFPE) tumor section^19, 51, 52^. Although a single HRP-antibody is generally applied for each work cycle, the use of different colored HRP-active fluorescent dyes permits visualization of 3 to 4 cell markers in a FFPE tumor section together with the HIAR process^53^. In addition, more cell markers can be localized by adding a H_2_O_2_-mediated fluorescence stripping step in the repeated ‘staining-imaging-stripping’ procedure^53^. H_2_O_2_-based fluorophore oxidation is simple and does not require any special antibody, fluorescent dye, or equipment for multiplex microscopy, but it is still not compatible with volumetric tissues. We have tested this approach in thick tumor tissues; due to the generation and trapping of oxygen bubbles in the middle of the tissues, H_2_O_2_ results in the disruption of tissue morphology and integrity (**Supplementary Figures 10**). In addition, repeated treatment of a tissue sample with oxidizing chemicals could damage protein antigens and prevent imaging of a number of cell markers^54, 55^. Therefore, it is necessary to develop a non-chemical-based fluorescence bleaching method for 3D multiplex IF imaging of volumetric tumor tissues.

Here, we introduce a LED-mediated photobleaching methodology for 3D multiplex IF microscopy of optically cleared tumor tissues. By integrating the LED-mediated photobleaching process with the T3 procedure, we built a cyclic workflow that allowed for imaging multiple cell markers in the same whole 400 µm-thick tumor macrosection. We previously reported the T3 method in which simple immersion of tissue macrosections in aqueous D-fructose solutions enabled the achievement of adequate tissue transparency for high-resolution 3D confocal fluorescence microscopy of whole tumor macrosections without noticeable changes in tissue volume and protein antigens^32, 33, 34^. Light from the LED system effectively penetrated and bleached fluorescence throughout a D-fructose-mediated optically cleared tumor macrosection. Photobleached tumor macrosections were then quickly prepared for the next cycle of IF staining by readily removing D-fructose *via* washing in PBS. We also implemented a framed glass slide-mediated novel tissue mounting and tissue region-tracking methods. DAPI-enabled repeat 3D microscopy allowed for imaging each tumor macrosection at the same X/Y/Z positions and reconstructing 3D multiplex images at tissue-wide and cellular resolution.

Previous studies have demonstrated that light irradiation using custom LED devices effectively reduced autofluorescence in thin FFPE tissue sections, providing IF and fluorescent *in situ* hybridization (ISH) image data with a high signal/noise ratio^56, 57^. However, use of an LED to strip the fluorescence signals in an IF-stained tissue remains untested. To the best of our knowledge, our work is the first study to develop a cyclic workflow involving LED-mediated photobleaching for 3D multiplex IF microscopy of optically cleared tissues.

We validated the workflow for 3D IF microscopy of cancer cells (CK8), vascular cells (αSMA, CD31, ER-TR7), and immune cells (CD8, CD45, CD3) in the same whole TUBO mouse mammary tumor macrosection by conducting three work cycles with DAPI and fluorescent primary antibodies. After reconstructing 3D tissue-wide and cellular resolution multiplex images, we performed quantitative 3D spatial analysis of the TME in terms of spatial cellular phenotyping, volume occupancy, and distribution patterns of different cell types, 3D structural features of cellular organization, and spatial cell-cell interactions. Based on those 3D spatial readouts, we demonstrated cell-level regional remodeling of the TME and higher tumor-infiltrating CD45^+^CD8^+^CD3^+^ CTLs after treatment with STING agonist, DMXAA. By assessing multiple components in a tumor simultaneously, our method provides a unique window into tumor heterogeneity and response to therapy.

Although our study demonstrated a new approach for increasing multiplexing power in 3D optical microscopy of cleared tumor tissues, we also recognize the need for shortening photobleaching time and developing a high-throughput tissue processing system. There are high-power LED (>100W) chips for building a multi-LED array that not only speeds up photobleaching but also carries out several tumor macrosections simultaneously. Our cyclic workflow can be also adapted to a microfluidic-based automated system for fast and consistent tissue processing^58, 59^ . We anticipate that these further developments would enable automated high-throughput 3D multiplex imaging of large volumetric tumor tissues. Taken together, the advanced multiplex imaging method we have presented generates multiscale 3D maps of diverse biomarkers throughout a large tumor tissue and offers comprehensive spatial information of the TME structure and tissue- and cell-level responses to a cancer therapy.

## Methods

### LED photobleaching system

100 W warm white LED chip (2800 - 3500mA, 30 - 34V DC, 8 - 9K Luminous Flux) and other color light LED chips (2800 - 3500mA, 20-34V DC, 3-8K Luminous Flux) were purchased (Chanzon Technology). A LED chip was assembled with an aluminum heatsink (80 x 67 mm (L x W)) and cooling fan (79 x 79 x 29 mm (L x W x H), 2V DC, 0.2A). After connecting with a power supplier (Input: AC 85-265V, 50/60Hz, Output: DC 20-38V, 3000mA, 100W) by soldering iron, the LED device was placed in a refrigerator (48 x 47 x 85 cm^3^ (W x D x H)) with temperature set at 4°C. To further control temperature during photobleaching, dry ice (0.5-1 kg) was added in the bottom shelf of the refrigerator. Screws were installed on the top surface of the heatsink as the holders of a tissue glass slide and then wrapped with parafilm tape on its head to prevent slipping of the glass slide. By adjusting the height of the individual screws, the distance between the bottom of tissue sample and the top of LED chip was set to 2.1 cm with the consideration of the thickness of a tissue glass slide (1 mm). To determine the temperature at the sample stage during photobleaching, a water-containing glass dish was placed on the LED device in the refrigerator with dry ice and the temperature of the water was measured at 0, 2, 4, 6, and 8 hrs after light irradiation using a thermometer. The emission spectra of the LED chips were measured using a photospectrometer (USB4000, Ocean Insight).

### Mouse tumor model and treatment with DMXAA

TUBO cells, derived from a spontaneous mammary tumor in a BALB-NeuT female mouse ^60^, were obtained from Dr. Steve Kron (University of Chicago) and cultured in Dulbecco’s Modified Eagle’s Medium-High glucose (SIGMA, Cat#: D5671) supplemented with 20% heat-inactivated FBS, and 1% penicillin in 5% CO_2_ and at 37°C. BALB/c female mice (6-8 weeks old) were purchased from Envigo. 2x10^5^ TUBO cells in 50 µl PBS were injected into the 4^th^ mammary fat pad of each mouse. TUBO tumors were developed and reached 5-8 mm in diameter at 10-14 days after injection of the cancer cells. DMXAA (10 μL at 1 mg/mL in PBS, MilliporeSigma, Cat#: D5817) was directly injected into a TUBO tumor in the mouse model using a 29G x 1/2” insulin syringe (Exel Comfort Point, Cat#: 26029). At 2 days post-treatment, the mouse was sacrificed, and the tumor was collected.

### Fluorescent antibodies

Primary monoclonal antibodies were purchased from commercial vendors; anti-CD31 (BioLegend, MEC13.3), anti-ER-TR7 (BioXcell, ER-TR7), anti-αSMA (MilliporeSigma, 1A4), anti-CD8 (BioXcell, 2.43), anti-CD45 (BioLegend, 30-F11), anti-CD3 (BioXcell, 17A2), and anti-CK8 (BioLegend, 1E8). Antibodies at concentration of 0.5-1 mg/mL in PBS (pH 8.0) reacted with amine-reactive fluorophores, including FITC (MilliporeSigma, Cat#: F7250), sulfo-Cy3-NHS ester (Lumiprobe, Cat#:11320), sulfo-Cy5-NHS ester (Lumiprobe, Cat#:13320), DyLight(DL)488-NHS ester (Thermo Scientific, Cat#: 46402), DL550-NHS ester (Thermo Scientific , Cat#: 62262) and DL633-NHS ester (Thermo Scientific, Cat#: 46414), at a molar ratio of 1:20 (antibody:fluorophore). Unconjugated, free fluorophore molecules were removed by dialysis purification using cassettes (Thermo Scientific, MWCO 10K) in 1 L PBS at 4°C for 3 days with daily PBS refreshment. Fluorescent antibody solutions were stored at 4°C in a dark environment.

### Tissue processing, immunofluorescence staining, and optical clearing

The protocols for tissue processing, IF staining, and optical clearing were adapted from our previous reports on transparent tissue tomography (T3) method^32, 33^. In brief, collected TUBO tumors were embedded in 2% agarose gel (dissolved in distilled water, LE Quick Dissolve Agarose, GeneMate) in 12-well plates. Then, the tumors were sectioned to 400 μm thickness using a vibratome (VT1200S, Leica) and macrosections were fixed in 2% paraformaldehyde (PFA) solution for 15 mins at room temperature followed by PBS washing. The agarose gel surrounding the tumor in each macrosection was cut with a razor blade to mark it for positioning and mounting on a microscope glass slide with a ‘L’ shaped frame for sample orientation on X and Y axis. For immunofluorescence staining, 0.7 µL of DAPI (at 5 µg/mL, Thermo Scientific, Cat#: 62248) and 5 µL of each fluorescent antibody (at 0.5 or 1 mg/mL) were mixed in a staining buffer composed of 0.5 mL RPMI1640 cell culture media (Gibco^TM^, Cat#: 22400089) with 1% (w/v) IgG-free bovine serum albumin (BSA, MilliporeSigma, Cat#: A2058). The tumor macrosection was incubated in the antibody cocktail solution for 18 hrs at 4°C under gentle shaking. After washing 3 times in PBS, stained tumor macrosections were incubated in a series of 10 mL D-fructose (MilliporeSigma, Cat#: F0127) solutions at different concentrations (20%, 50%, 80%, 100% (w/v) in 10 mM phosphate buffer (pH 7.8)), each for 30 mins at 4°C under gentle agitation. For additional IF staining after the photobleaching step, the macrosections were subjected to a reverse tissue clearing process of PBS washing followed by repeat staining with the next set of fluorescent antibodies.

### LED-mediated fluorescence bleaching

An IF-stained, cleared tumor macrosection was mounted on a microscope glass slide with an ‘L’ shaped frame made of plastic tape. A glass coverslip was placed on the top of the macrosection to prepare a ‘sandwich’ mount for upright confocal microscopy (**Supplementary Figure 3A**). After microscopy, the top glass coverslip was removed and the tumor macrosection on the glass slide put on the sample stage of the LED device. Additional fructose solution drops were added to the tumor macrosection to prevent sample drying. The LED light was turned on by plugging the power supplier in and the glass window of the refrigerator was covered with a paper sheet during photobleaching.

### 3D confocal fluorescence microscopy

3D fluorescence images were acquired using an upright confocal microscope (Caliber ID, RS-G4), 20X/0.8 NA dry objective (Olympus, UPLXAPO20X), and 40X/1.4 NA oil objective (Olympus, UPLXAPO40XO). 3D scanning of macrosections was performed at defined voxel (X×Y×Z) sizes (0.614×0.614×3.05 μm^3^ and 0.313×0.313×1.52 μm^3^ for tissue-wide and cellular resolution 3D images, respectively) with 8 frame averaging using the following wavelengths: 405 nm excitation laser and 415-485 nm emission filter for DAPI; 488 nm excitation laser and 498-542 nm and 506-594 nm emission filters, respectively, for FITC and DL488; 561 nm excitation laser and 574-626 nm emission filter for Cy3 and DL550; 640 nm excitation laser and 662-738 nm emission filter for Cy5 and DL633; and 785 nm excitation laser and an open emission filter for tissue reflection. Raw 16-bit 3D image data were saved to a hard drive connected to a data storage server.

### Construction and processing of 3D multiplex tumor images

Open-source plugins on Fiji software, ‘BigWarp’ and ‘Fijiyama’, have been used for constructing 3D multiplex tumor images. Individual 3D IF images of different cell markers in a whole tumor macrosection were aligned and registered into a multiplex 3D image by manually assigning areas matched between the images from different work cycles and then performing ‘Thin Plate Spline’ transformation in the BigWarp plugin^61^. High-resolution 3D multiplex images of selected regions in a tumor macrosection were constructed by transforming and aligning individual fluorescence channel images using ‘training’ mode and ‘automatic registration’ function in the Fijiyama plugin^62^. 3D renderings of multiplex image data were created using Imaris software version 10.0 (Oxford Instrument).

For quantitative measurement of LED-mediated fluorescence bleaching (**Figure 2** **and Supplementary Figure 2, 4**), we masked a tumor region in 16-bit 488, 561, and 633 channel images of a tumor macrosection and determined the total area intensities using Fiji (ImageJ) software. The fluorescence intensity ratio was calculated by *R* = (*I*_i_ – *I*_b_) / (*I*_s_ – *I*_b_) where *I*_b_, *I*_s_, *I*_i_ represent the total intensities of the masked same tumor area in the channel images obtained pre-IF staining (tissue autofluorescence background), post-IF staining but prior to photobleaching, and different time points after starting photobleaching, respectively. Intensity ratios of ‘0’ and ‘1’ indicate tissue autofluorescence background and full fluorescence intensity prior to photobleaching, respectively.

To measure volume % of caner and vascular cell types in the control and DMXAA-treated tumor macrosections (**Figure 4B**), we first performed hyperstack segmentation of DAPI^+^, CK8^+^, αSMA^+^, CD31^+^, and ER-TR7^+^ cell areas in 3D whole macrosection images with individual cutoff thresholds in which all pixels in a raw channel image corresponded to each cell or nucleus marker using Fiji software. Then, we converted the raw 3D images to 8-bit binary hyperstack images, measured ‘area’ value in each stack of images, and obtained ‘total area’ value by sum. The volume % of each cell type was calculated by *V %* = (Total area of a cell type / Total area of DAPI^+^ tumor parenchyma) × 100.

To profile the radius of ER-TR7^+^ segments from the control and DMXAA-treated tumor macrosections (**Figure 4C**), we opened the previously acquired 8-bit binary hyperstack images of ER-TR7^+^ segments in Amira 3D Pro software (Ver.2023.1.1) and processed ‘Skeletonization’ to define the centerlines of ER-TR7^+^ stroma branches using ‘Auto skeletonization’ Amira module based on TEASAR (tree-structure extraction algorithm for accurate and robust skeletons)^63^. The radius and number (in *log_10_* #) of ER-TR7^+^ segments were computed based on the defined centerlines through the fibers. 3D pseudo-color images according to the radius were also rendered (**Supplementary Figure 6**).

For quantitative 3D spatial analyses of tumor region images, we segmented CK8^+^, ER-TR7^+^, CD8^+^, CD45^+^, and CD3^+^ cells in high-resolution hyperstack images using Imaris software and its ‘Surface’ modules. The cutoff thresholds were determined by interactive comparison of signals in the raw image data. We used ‘Split touching Objects (Region Growing)’ module for accurate and fully representative individual cell surface segmentation. We further processed the segmented images with ‘Overlapped Volume’ filter as inclusive or exclusive criteria for quantitative determination of immune cell subtype populations and cancer-immune cell interaction.

The volume % of a CD45^+^ immune cell subtype in region #1 (**Figure 5B**) was calculated by *V* % = (Volume of immune cell segments / Volume of region) × 100. The volume % of CK8^+^ tumor cells interacted with CTLs in region #1(**Figue 5D**) was calculated by *V* % = (Volume of CK8^+^ tumor cell segments bounded to CTL segments / Volume of total CK8^+^ tumor cell segments) × 100.

### Statistical analysis

Values are displayed as mean ± standard deviation (SD), and statistics were run using multiple unpaired t-tests in Prism software. If the *P* value is less than 0.05, it is considered as “significant.”

## Acknowledgements

We thank Drs. Stephen Kron and Vytautas Bindokas at the University of Chicago for providing the TUBO cell line and consulting on image processing and analysis. We also thank Drs. Xincheng Yao and Taeyoon Son at University of Illinois Chicago for helping us measure emission spectra of LED chips. This work was supported by the National Institute of General Medical Sciences R35 GM142743 (to S.S.-Y.L.) and University of Illinois Cancer Center Pilot Project Awards 2020-PP-07 and 2023-29-UICCPG (to S.S.-Y.L.)

## Author contributions

J.Z. and S.S.-Y.L. conceived the concept and designed the study. X.W., J.Z., and S.S.-Y.L. built and tested the LED photobleaching system. Y.-C.W. and J.Z. prepared, treated and processed mouse tumor tissues. J.Z. performed immunofluorescence staining and confocal microscopy. E.H.P. and J.Z. conducted 3D image data processing and quantitative analyses. J.Z., E.H.P. and S.S.-Y.L. wrote and edited the manuscript. All authors approved the final manuscript.

