## Supplementarydata for "Increased multiplexity in optical tissue clearing-based 3D immunofluorescence microscopy of the tumor microenvironment by LED photobleaching"

**Supplementary Figure 1**

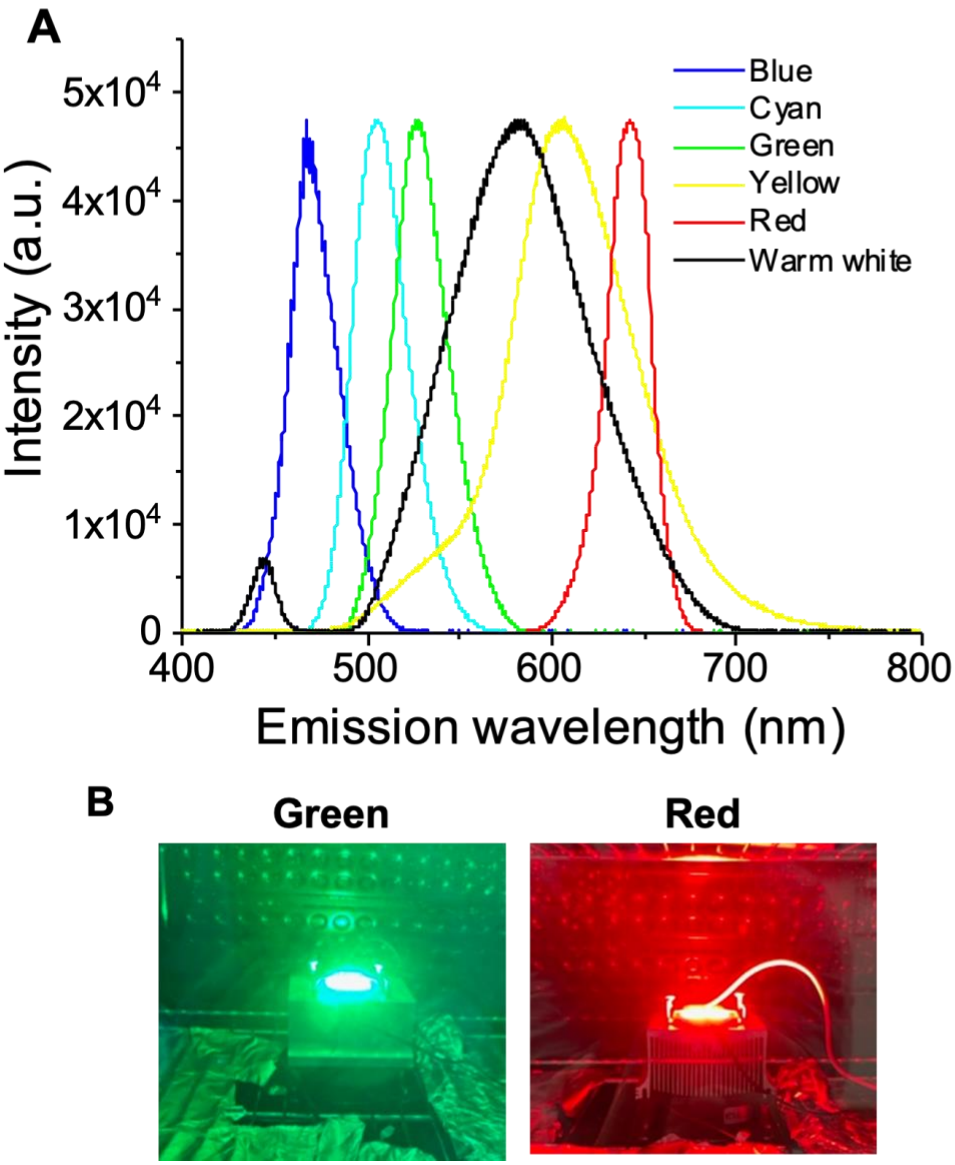


**Supplementary Figure 1. High-power color light LED systems. (A)** Emission spectra of 100W LED chips lighting in different colors. **(B)** Photographs of green and red light LED irradiation systems.

#### Supplementary Figure 2

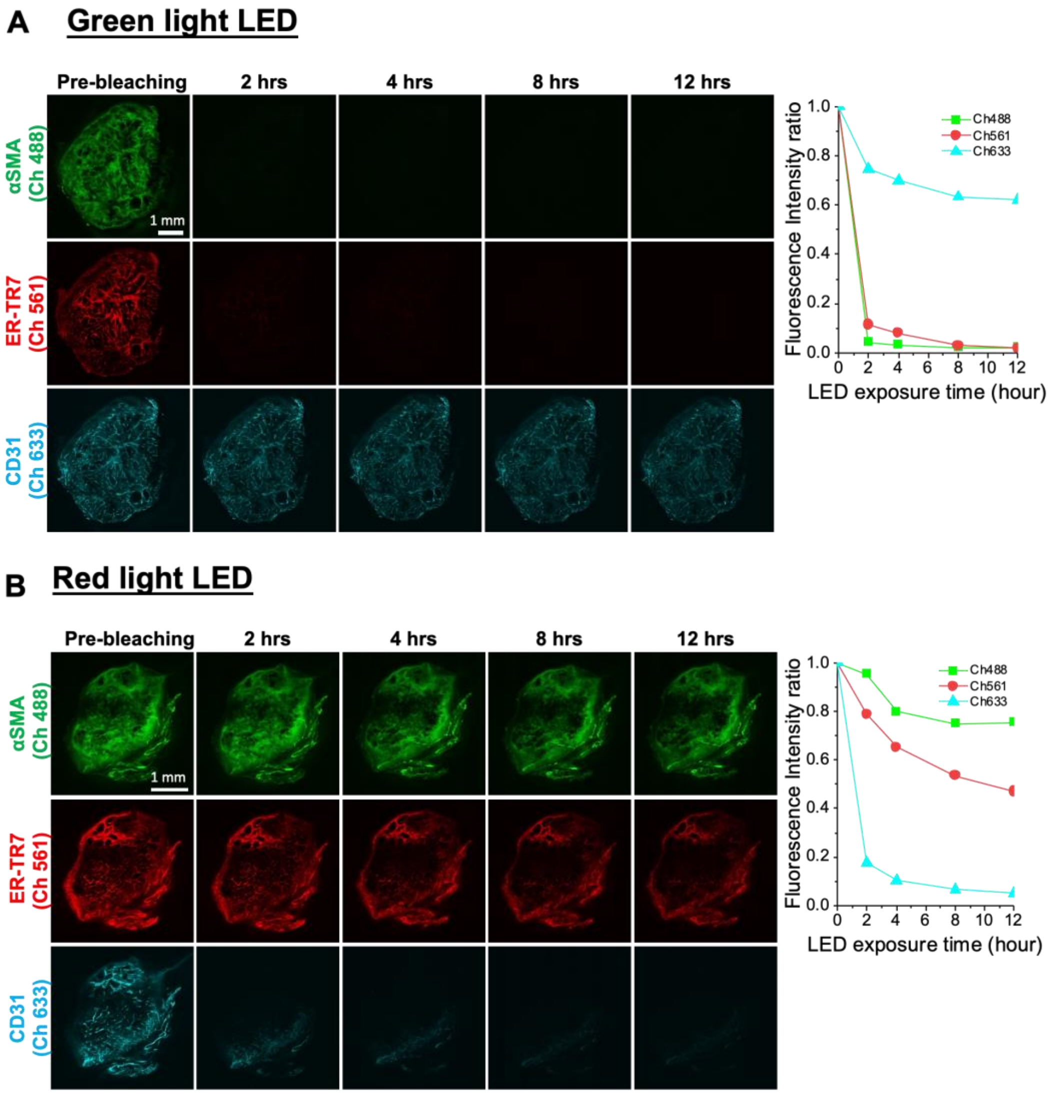


**Supplementary Figure 2. Selective fluorescence bleaching using color light LED irradiation systems. (A)** Mosaic fluorescence channel images of αSMA (green, excitation at 488 nm), ER-TR7 (red, excitation at 561 nm), CD31 (cyan, excitation at 633 nm) on the LED-far side (top surface) of 400 µm-thick tumor macrosections before and after exposure to the LED green (top) and red (bottom) light for 2, 4, 8, and 12 hours. Scale bar: 1 mm. **(B)** Quantification of fluorescence intensity changes in 488, 561, and 633 channel images from the top surfaces of the tumor macrosections during green and red light LED-mediated photobleaching. The intensity ratio was plotted based on the pre-IF staining ‘0’ and pre-bleaching ‘1’ levels.

**Supplementary Figure 3**

#### A X-Y axis alignment

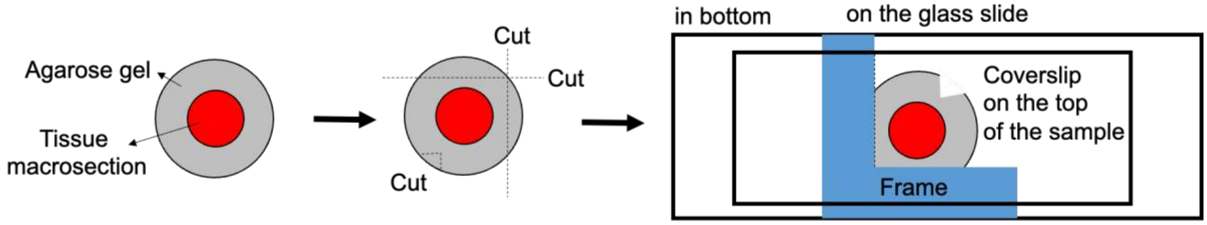


#### B Z axis alignment

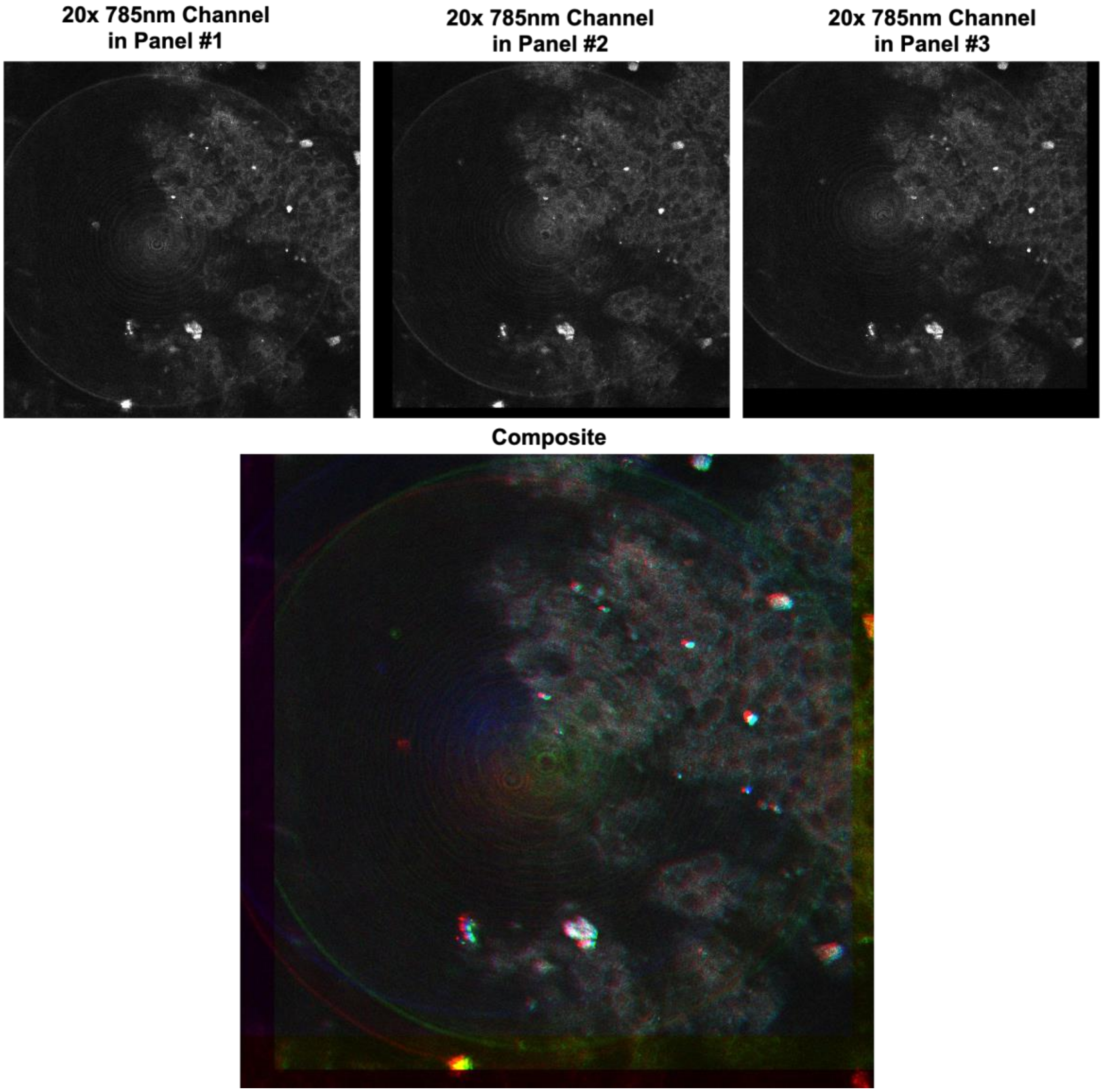

**Supplementary Figure 3. Alignment of X/Y/Z position of a tissue sample and microscope imaging for repeat 3D immunofluorescence of a tumor macrosection. (A)** X-Y axis alignment by positioning a tissue sample. The agarose gel surrounding a tumor macrosection is cut to be fitted flush to the 'L' frame on a glass slide. A coverslip is put on the top of a tumor sample. **(B)** Z axis alignment by microscope scanning at the same top (Z) position of the tissue sample. 785 nm excitation (with open emission filter)-mediated tissue reflection images show the same Z scanning position of a tumor macrosection for 3D imaging of cell marker panel at each work cycle. The 785 nm channel images obtained from three work cycles were merged into a composite image (bottom).

#### Supplementary Figure 4

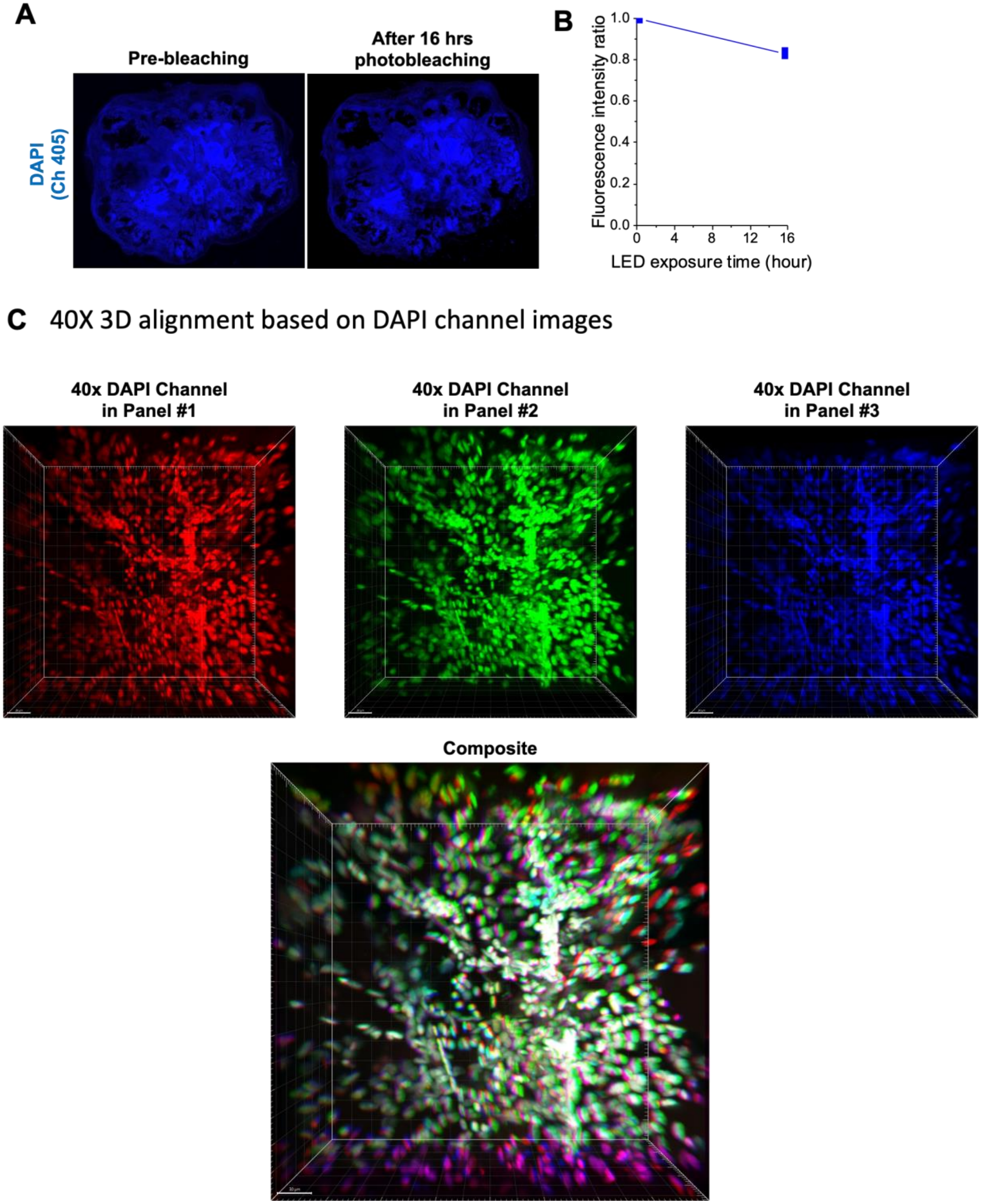

**Supplementary Figure 4. Tracking DAPI signal and pattern for repeated high-resolution 3D microscopy of selected regions in a tumor macrosection. (A)** Mosaic images of DAPI signal (excitation at 405 nm) in a tumor macrosection before and after 16 hours of photobleaching using the warm white LED irradiation system. **(B)** Quantification of fluorescence intensity change in the DAPI channel images before and after 16 hours photobleaching. The intensity ratio was plotted based on the pre-staining ‘0’ and pre-bleaching ‘1’ levels. **(C)** Repeated high-resolution 3D DAPI channel images of a selected region in a tumor macrosection produced through three work cycles. The DAPI channel images (pseudo-colored in red, green, blue) from three work cycles were registered into a composite image (bottom) using Fijiyama plugin. White color in the composite image indicates the complete registration of DAPI signal patterns in individual high-resolution 3D images from different work cycles.

### Supplementary Figure 5

#### Control tumor

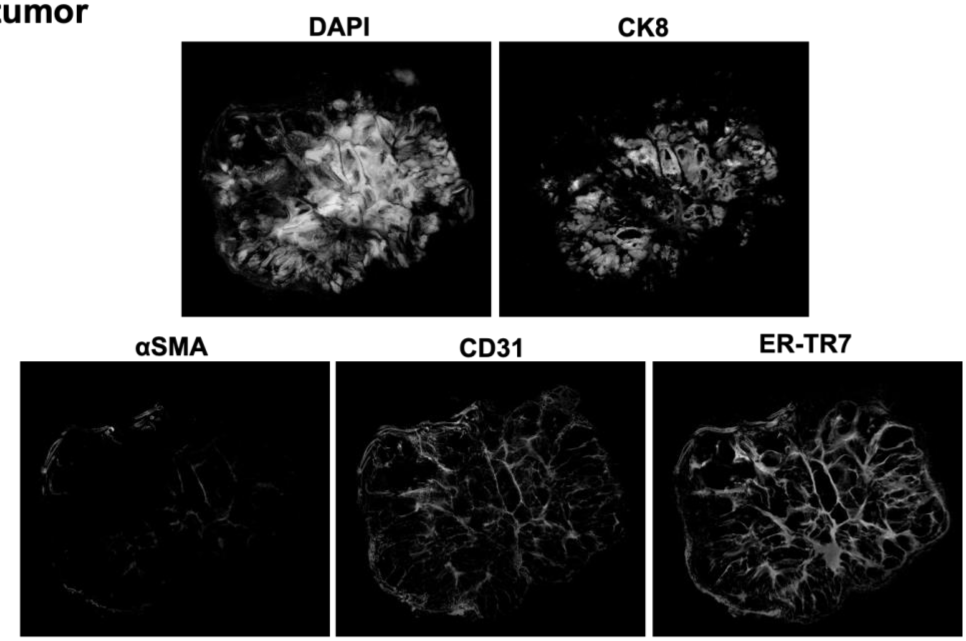


#### Treated tumor with DMXAA

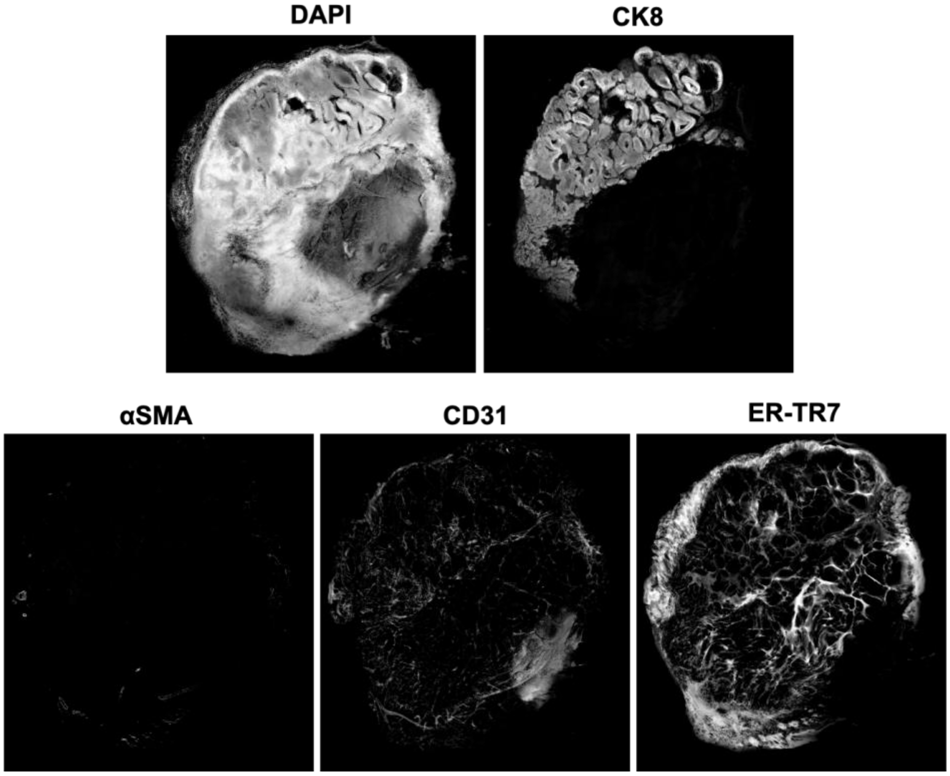

**Supplementary Figure 5. Segmentation of DAPI$^+$ tumor parenchyma, CK8$^+$ tumor cell, αSMA$^+$, CD31$^+$ and ER-TR7$^+$ vascular cell markers in 3D images of the whole control and treated tumor macrosections.** 8-bit binary images of the segments shown by maximum intensity projections using Fiji.

#### Supplementary Figure 6

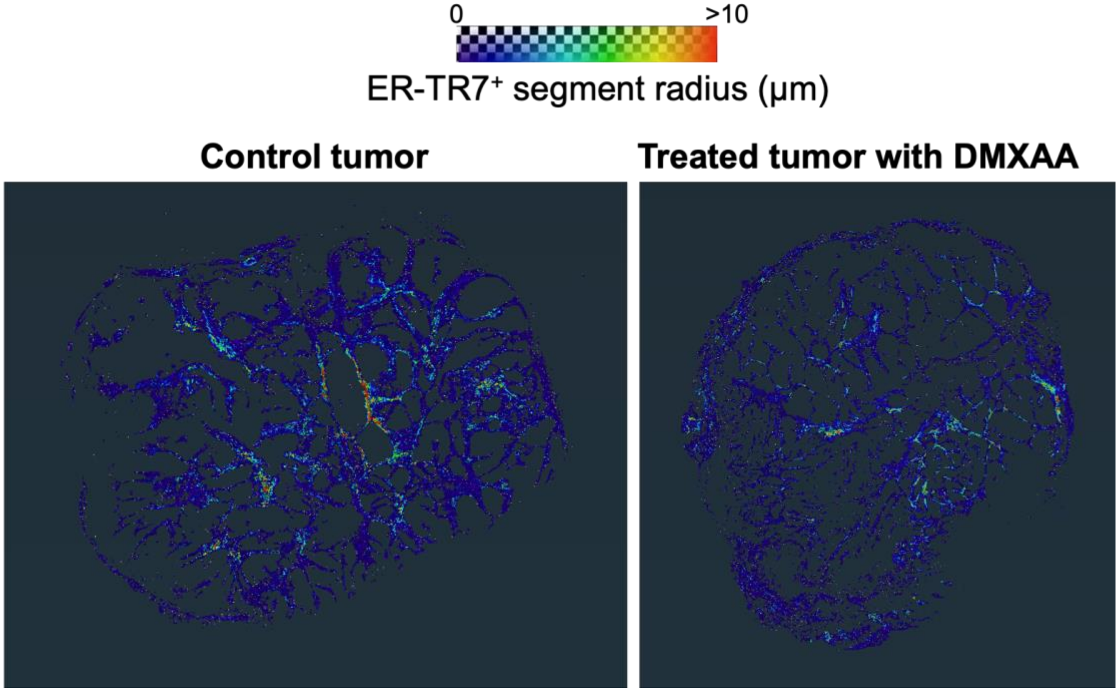


**Supplementary Figure 6. 3D pseudo-color mapping of ER-TR7$^+$ fibrotic stroma according to branch thickness.** The color scale corresponds to the average radius of each identified ER-TR7$^+$ segment. Red: largest (>10 μm).

**Supplementary Figure 7**

**Control tumor**

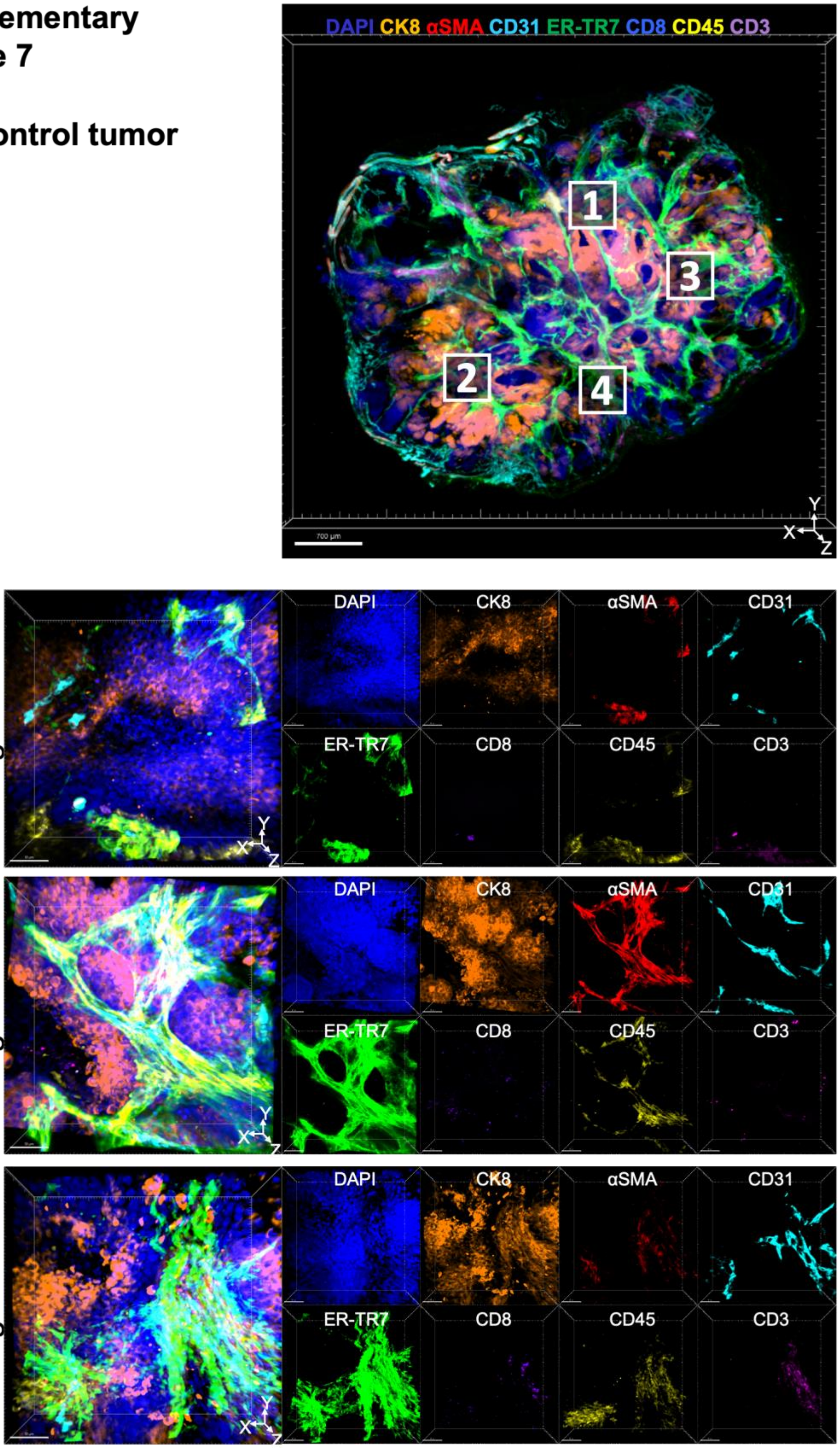

**Supplementary Figure 7. High-resolution 3D multiplex microscopy of selected regions in the control tumor macrosection.** Other than Region #1 **(Figure 4)**, an additional three regions (Region #2-4, white solid boxes in the top image of the whole tumor macrosection) were selected and imaged in 3D.

**Supplementary Figure 8**

**Treated tumor with DMXAA**

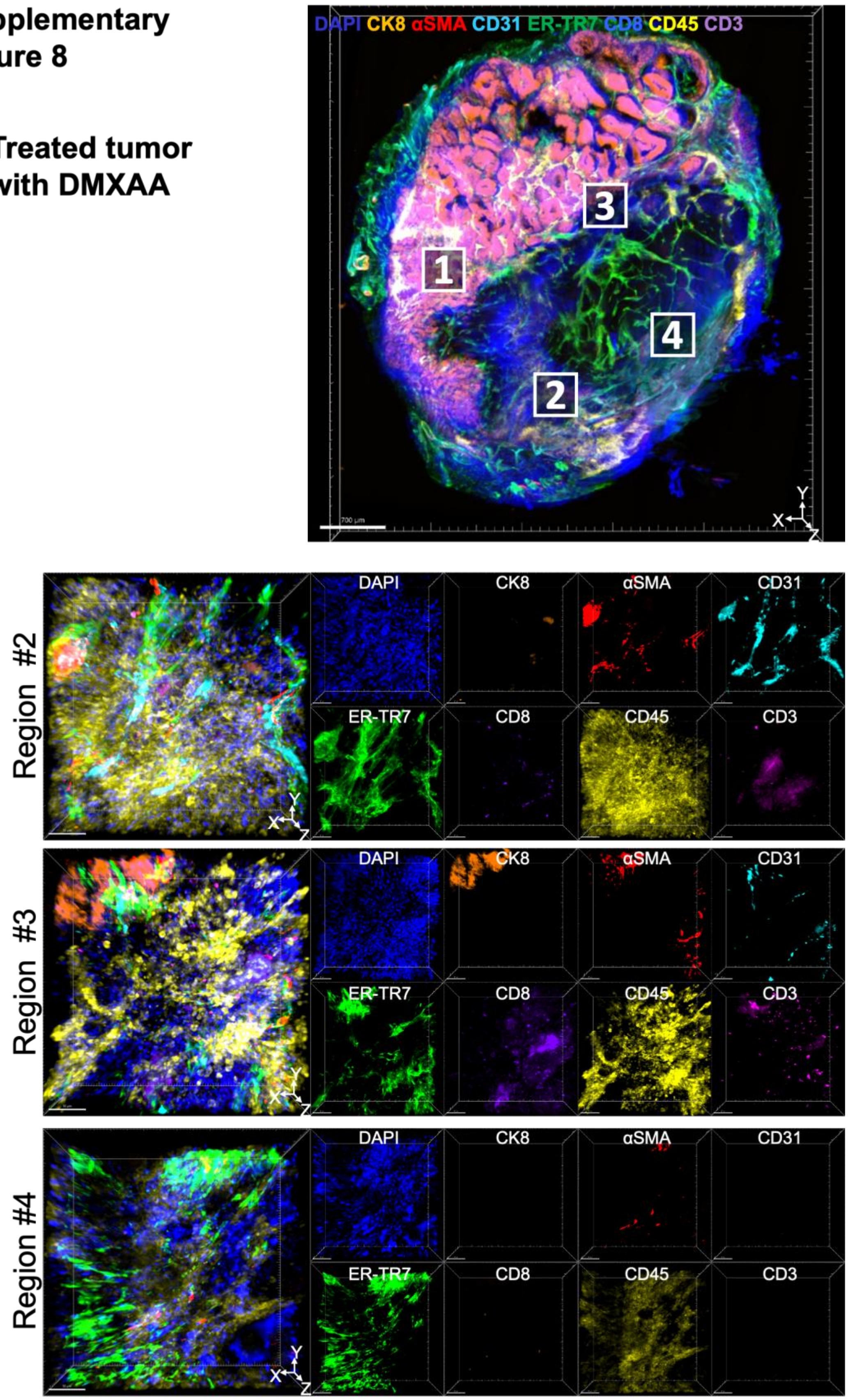

**Supplementary Figure 8. High-resolution 3D multiplex microscopy of selected regions in the treated tumor macrosection.** Other than Region #1 **(Figure 4)**, an additional three regions (Region #2-4, white solid boxes in the top image of the whole tumor macrosection) were selected and imaged in 3D.

#### Supplementary Figure 9

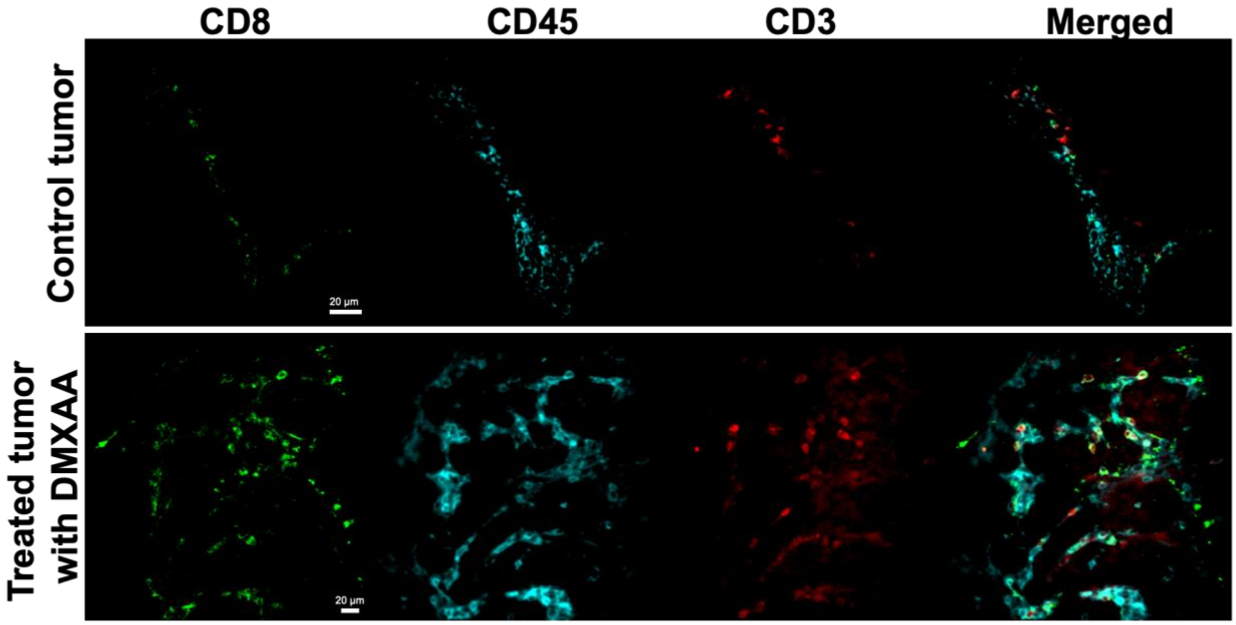


**Supplementary Figure 9. 2D images of immune cell markers (CD8, CD45, CD3) in the middle of high-resolution 3D images of Region #1 in the control and treated tumor macrosections**. The images show IF-stained CD markers on the membranes of individual immune cells. Scale bar: 20 μm.

### Supplementary Figure 10

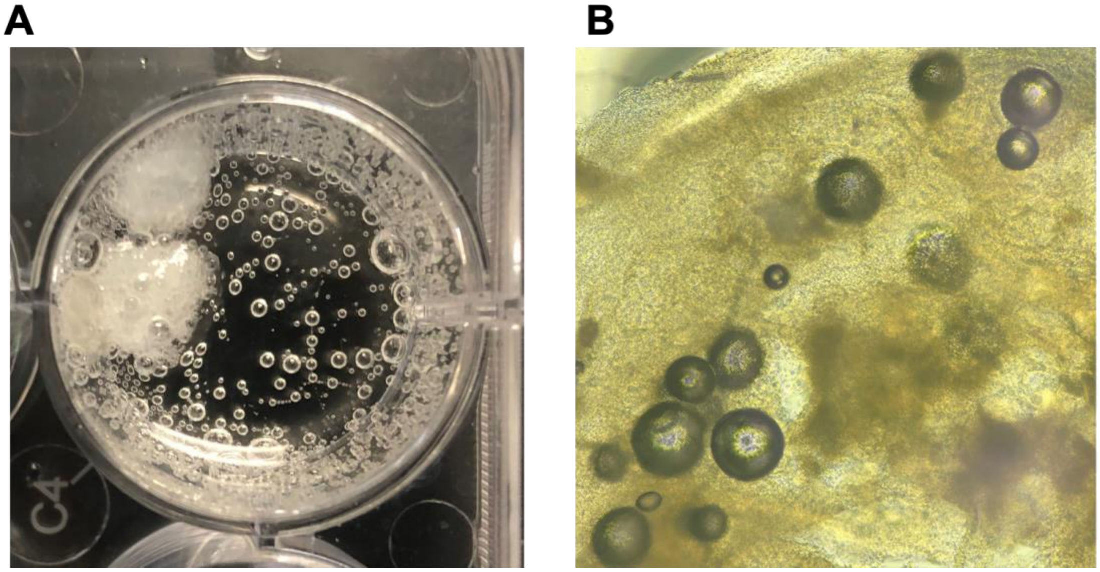


**Supplementary Figure 10. Oxygen bubbling and bubble trapping in tumor macrosections during and after incubation in $H_2O_2$ solution. (A)** Photograph of tumor macrosections in 3% $H_2O_2$ solution (in PBS, pH=10). **(B)** Light microscope image of oxygen bubbles within the tumor macrosection treated with $H_2O_2$ solution.

**Supplementary Video 1. 3D rendering of 7-plex tumor macrosection image.**

**Supplementary Video 2. 3D rendering of high-resolution Region #1 image from the control tumor macrosection.**

**Supplementary Video 3. 3D rendering of high-resolution Region #1 image from the treated tumor macrosection.**
